# Museum epigenomics: characterizing cytosine methylation in historic museum specimens

**DOI:** 10.1101/620583

**Authors:** Tricia L. Rubi, L. Lacey Knowles, Ben Dantzer

**Author notes:** Correspondence: Tricia L. Rubi. Department of Biology, University of Victoria, 3800 Finnerty Rd, Victoria, BC, V8P 5C2, Canada.

## Abstract

Museum genomics has transformed the field of collections-based research, opening up a range of new research directions for paleontological specimens as well as natural history specimens collected over the past few centuries. Recent work demonstrates that it is possible to characterize epigenetic markers such as DNA methylation in well-preserved ancient tissues. This approach has not yet been tested in traditionally-prepared natural history specimens such as dried bones and skins, the most common specimen types in vertebrate collections. In this study, we develop and test methods to characterize cytosine methylation in dried skulls up to 76 years old. Using a combination of ddRAD and bisulfite treatment, we characterized patterns of cytosine methylation in two species of deer mouse (*Peromyscus spp.*) collected in the same region in Michigan in 1940, 2003, and 2013-2016. We successfully estimated methylation in specimens of all age groups, though older specimens yielded less data and showed greater interindividual variation in data yield than newer specimens. Global methylation estimates were reduced in the oldest specimens (76 years old) relative to the newest specimens (1-3 years old), which may reflect *post mortem* hydrolytic deamination. Methylation was reduced in promoter regions relative to gene bodies and showed greater bimodality in autosomes relative to female X chromosomes, consistent with expectations for methylation in mammalian somatic cells. Our work demonstrates the utility of historic specimens for methylation analyses, as with genomic analyses; however, such studies will need to accommodate the large variance in the quantity of data produced by older specimens.

---

Museum collections worldwide house billions of specimens and are an invaluable resource for tracking how organisms change over time. One of the most influential fields in modern collections-based research is museum genomics, which is transforming the way that museum specimens are used in research by enabling studies of long term change in genetic variation. Until recently, museum genomics research focused exclusively on genetic sequences; however, a growing body of recent work in “paleoepigenetics” demonstrates that ancient DNA retains patterns of *in vivo* DNA methylation (Orlando and Cooper 2014; Gokhman et al. 2016), a well-studied epigenetic mechanism associated with transcriptional regulation and modulation of gene expression (Jones 2012). The implications of this discovery are compelling; methylation markers in museum specimens could elucidate patterns of gene expression in past populations, opening up a number of new directions for collections-based research. In addition, the ability to document how epigenetic effects change over time may help clarify the role of epigenetic processes in adaptation and evolution.

Around a dozen paleoepigenetic studies have been published to date (Briggs et al. 2010; Llamas et al. 2012; Gokhman et al. 2014; Pedersen et al. 2014; Smith et al. 2014, 2015; Orlando and Cooper 2014; Seguin-Orlando et al. 2015; Gokhman et al. 2016; Hanghøj et al. 2016; Gokhman et al. 2017; Murphy and Benítez-Burraco 2018). To our knowledge, all previous studies have focused on ancient DNA from paleontological and archaeological specimens rather than “historic DNA” from museum specimens collected by naturalists in the modern era, which range from decades old to a few centuries old.

Historic specimens are more abundant and broadly available across taxa than ancient specimens, and can therefore be used for a greater diversity of study questions. Though researchers now routinely collect tissue vouchers for genomic analyses, traditional preparations such as dried skins and bones still comprise the majority of existing vertebrate collections and represent some of the oldest and rarest specimens. Somewhat counterintuitively, such historic tissues are not necessarily more amenable to genomic work than ancient (*i.e.*, paleo) tissues. Historic specimens have the advantage of being much “younger” than paleontological specimens, reducing the amount of time for *post mortem* DNA damage to accumulate. Such specimens are also likely to be more pristine, harboring less exogenous DNA, and have been stored in (hopefully) optimal conditions. However, high quality ancient specimens such as tissues obtained from permafrost are often remarkably well-preserved and may actually be less degraded than historic bones and skins despite their age. DNA degradation such as fragmentation and nucleotide damage (notably hydrolytic deamination) is the primary challenge for ancient and historic DNA studies, making DNA harder to extract and amplify, increasing contamination risk, and producing sequence errors due to base pair misincorporations (Willerslev and Cooper 2005). Nevertheless, the field of museum genomics is thriving, and new protocols and analytical methods continue to broaden and strengthen collections-based genomic analyses. Llamas et al. (2012) remark that the main challenge in ancient methylation protocols is extracting amplifiable nuclear DNA, which is now feasible even for low quality historic specimens such as bones and dried skins (*e.g.*, Irestedt et al. 2006; Bi et al. 2013).

In this study, we describe DNA methylation in skull specimens from deer mice (*Peromyscus spp.*) sampled from the same region in Michigan over three time periods: 1940, 2003, and 2013-2016. We generate reduced representation methylomes at base-pair resolution using a combination of double digest restriction site-associated DNA sequencing (ddRAD) and bisulfite treatment. To explore the effect of specimen age, we compare data yield and global methylation estimates in older versus newer specimens. For one of our species, we use genome annotations to describe methylation patterns in known genomic regions (putative promoters versus gene bodies and autosomes versus sex chromosomes). We conclude with a discussion of the challenges of working with historic samples, in particular loss of data, and the sampling designs and epigenetic analyses that can accommodate these challenges. We also highlight how epigenetic datasets, including the dataset produced in this study, can be used in future work to infer gene expression in past populations and characterize change over time in epigenetic effects.

## Methods

### Specimens and sampling design

We sampled 75 specimens total: 40 white-footed mice (*Peromyscus leucopus noveboracensis*) and 35 woodland deer mice (*Peromyscus maniculatus gracilis*). All specimens were collected from the same locality in Menominee county in Michigan over three collecting periods: 1940, 2003, and 2013-2016 (Figure 1). The specimens were traditional museum skull preparations (dried skulls stored at room temperature). When possible, we balanced sampling between the sexes. Skulls collected from 2013-2016 were provided by the Dantzer Lab at the University of Michigan and the Hoffman Lab at Miami University. Older skulls (1940-2003) were provided by the University of Michigan Museum of Zoology. Detailed specimen information is included in Supplementary Table S2.

**Figure 1:**
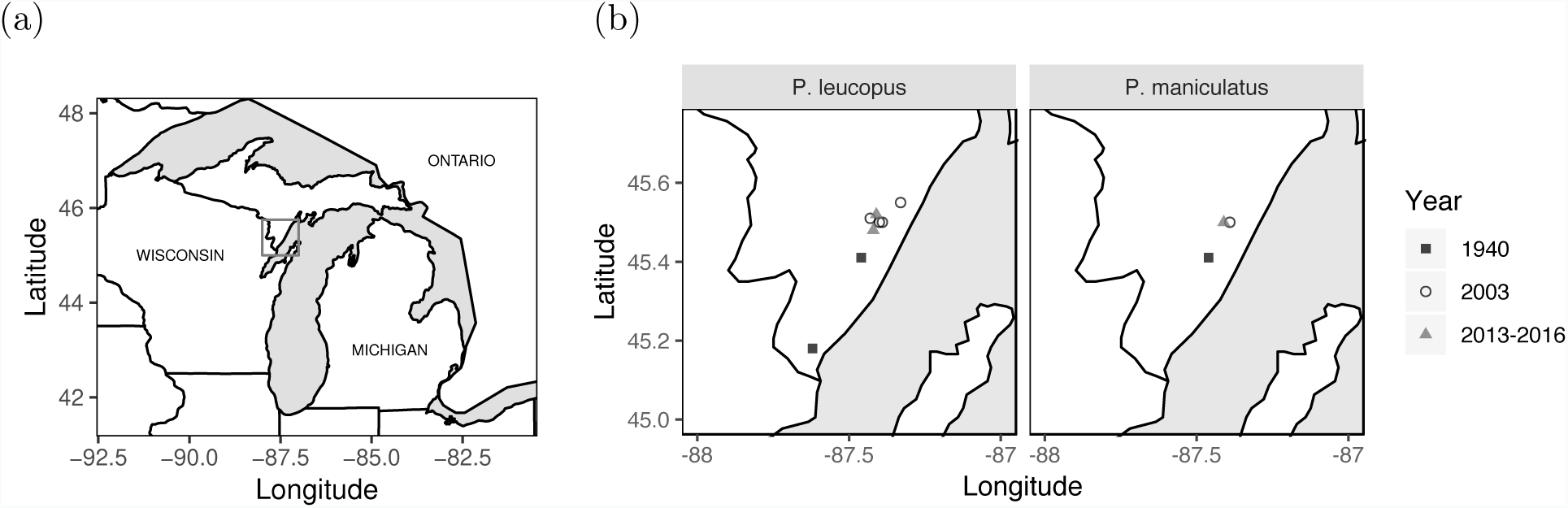
Sampled localities in the Upper Peninsula of Michigan. **(a)** The Great Lakes region of North America. The gray box indicates the region shown in (b). **(b)** Sampled localities for both species. Black squares indicate sampling in 1940, white circles indicate sampling in 2003, and gray triangles indicate sampling in 2013-2016.

### Tissue sampling and DNA extraction

All pre-amplification steps were performed in the ancient DNA facility in the Genomic Diversity Lab at the University of Michigan following standard protocols for working with historic DNA. Briefly, all work was performed under a hood in a dedicated laboratory for processing historic specimens and followed stringent anti-contamination protocols, including dedicated reagents, unidirectional flow of equipment and personnel, filtered pipette tips, and additional negative controls. We sampled tissue from traditional skull preparations (dried skulls stored at room temperature). To minimize damage to the skulls, we sampled microturbinates (small nasal bones) by inserting a sterile micropick into the nasal cavity to dislodge 5-12 mg of tissue (Wisely et al. 2004; Taylor and Hoffman 2010). Prior to DNA extraction, the bone fragments were placed into thick-walled 2 ml microcentrifuge tubes with four 2.4 mm stainless steel beads and processed in a FastPrep tissue homogenizer (MP Biomedicals) for 1 min at 6.0 m/s. All 2013-2016 specimens and some 2002-2003 specimens were extracted using a Qiagen DNeasy Blood and Tissue Kit with modifications for working with museum specimens. To increase yield, the rest of the specimens were extracted using a phenol-chloroform protocol. Detailed extraction protocols are described in the Supplementary Methods (also see Iudica et al. 2001; Mullen and Hoekstra 2008; Rowe et al. 2011).

### Library preparation

The samples were prepared for sequencing using a combination of double digest restriction site-associated DNA sequencing (ddRAD) and bisulfite treatment (see flowchart in Supplementary Figure S1, Supplementary Methods; also see Trucchi et al. 2016; van Gurp et al. 2016 for similar approaches). Samples were individually barcoded using a combinatorial indexing system (10 unique barcodes on the forward adapter and 10 unique indices on the reverse PCR primer) and processed into multiplexed libraries (see Supplementary Table S1 for oligonucleotide sequences). Specimens were assigned to the libraries based on the amount of DNA that could be extracted or specimen availability. We prepared three libraries with different starting concentrations of DNA - one high DNA concentration library (350 ng/specimen) of younger specimens (0-3 years old (yo)), one medium DNA concentration library (150 ng) of younger and older specimens (0-76 yo), and one low DNA concentration library (40 ng) of older specimens (13-76 yo). Two specimens were sequenced in both the medium and low concentration libraries.

We followed the ddRAD protocol outlined in Peterson et al. (2012) with added steps for bisulfite treatment. Briefly, we digested each sample with the restriction enzymes SphI-HF and MluCI for 1 hour at 37°C (New England Biolabs). These enzymes were chosen because they are insensitive to DNA methylation (and therefore will not show biased template enrichment) and have previously been used to prepare libraries in *Peromyscus* (Munshi-South et al. 2016). We added a spike-in of digested unmethylated lambda phage DNA (Sigma Aldrich) to each sample at a concentration of 0.1% of the sample concentration; these phage reads were used to directly measure the bisulfite conversion rate for each individual sample. We ligated custom methylated barcoded Illumina adapters (Sigma Aldrich) onto the digested products and pooled samples into sublibraries. Size selection was performed on a Pippin Prep electrophoresis platform (Sage Biosciences), with 376-412 bp and 325-425 bp fragments selected in the high and lower concentration libraries, respectively (a wider range was chosen for the latter to ensure that the samples exceeded the recommended minimum mass of DNA for the Pippin Prep cassette). Based on *in silico* digestion of the genomes, the estimated sampling rate for the selected restriction enzymes and size selection window was *c.* 25,000 loci. Bisulfite conversion was performed on the size selected sublibraries using a Promega MethylEdge Bisulfite Conversion Kit, which converted unmethylated cytosines to uracils, and amplified by PCR using KAPA HiFi HotStart Uracil+ MasterMix, which replaced uracils with thymines in the amplified product. Due to low DNA concentration in the final libraries for sequencing, the low concentration and medium concentration libraries were combined and sequenced on the same lane. The high concentration library was sequenced in one lane for 100 bp paired-end reads and the medium / low concentration library in a separate lane for 125 bp paired-end reads on an Illumina HiSeq 2500 (San Diego, CA).

### Illumina data processing

The raw sequence reads were demultiplexed using the *process_radtags* script of *Stacks* v.1.45 (Catchen et al. 2013) with a maximum allowed barcode distance of one (--barcode_dist 1). The restriction site check was disabled because bisulfite treatment can change the sequence at the restriction site (--disable_rad_check). Demultiplexed reads were trimmed for quality and adapter contamination and cut site sequences were removed using *TrimGalore* v.0.6.0 (www.bioinformatics.babraham.ac.uk/projects/trim_galore/). Quality and adapter trimming was performed using default settings for paired-end reads; by default, *TrimGalore* removes base calls with Phred ≤ 20, trims adapter sequences from the 3’ end, and removes sequences trimmed to a total length of 20 bp or less. The stringency for adapter trimming was set at the default minimum of 1 bp of overlap between the read sequence and adapter sequence; this highly stringent setting is recommended for bisulfite analyses because adapter contamination can skew methylation calling. After quality and adapter trimming, the reads were visually assessed for degradation at read ends using Mbias plots (Supplementary Figure S2). Cut site sequences were removed by trimming 5 positions from the 5’ end of forward reads (–clip_r1 5) and 4 positions from the 5’ end of reverse reads (–clip_r2 4). Forward reads were further trimmed to remove low quality positions at the read ends by trimming 5 more positions from the 5’ end and truncating reads to 118 bp at the 3’ end (–hardtrim5 118).

### Methylation calling

We focused on CpG methylation; in eukaryotes methylation almost always occurs on a cytosine, and in mammals almost exclusively in the context of a CpG dinucleotide (Jones and Takai 2001). Because methylation is tissue-specific, it is necessary to standardize the tissue sampled. We chose to sample bone tissue from dried skulls, one of the most common specimen types available in vertebrate collections. Even within a tissue the methylation state of a given CpG position in the genome may vary between alleles or across cells, so methylation at a given position is typically expressed as a percentage ranging from fully methylated (methylated in 100% of sequences) to fully unmethylated (methylated in 0% of sequences). Within a tissue, most CpGs are either fully methylated or fully unmethylated (though partial methylation is not uncommon), resulting in a bimodal distribution across loci (Rakyan et al. 2004; Eckhardt et al. 2006).

Paired-end reads were aligned to the appropriate genome (*Peromyscus maniculatus* NCBI ID: GCA_003704035.1; *Peromyscus leucopus* NCBI ID: GCA_004664715.1) and methylation calling was performed using the bisulfite aligner *Bismark* v.0.18.1 (Krueger and Andrews 2011). *Bismark* was run with default settings except for the mismatch criteria (-N 1) and gap penalties (--score_min L,0,-0.4), which were adjusted to allow more differences between the aligned reads and the reference. An analysis was also run with the default settings for both species and returned the same global methylation trends, but fewer loci; therefore, the results from the less stringent criteria are reported here. *Bowtie2* v.2.1.0 (Langmead and Salzberg 2012) was used as the core aligner. We also aligned the reads to the lambda phage genome (NCBI ID: NC_001416) using default alignment settings and used these reads to estimate the bisulfite conversion rate for each sample.

The methylation calls output by Bismark were further filtered for significance based on the sample-specific bisulfite conversion rate using functions from *MethylExtract* v.1.9 (Barturen et al. 2013). Briefly, we used Bismark to generate a list of all CpG positions in our sequences with the number of methylated and unmethylated reads. We then estimated the sample-specific bisulfite conversion rates from the lambda phage-aligned reads using the MethylExtractBSCR function. Significant methylation calls were determined using the *MethylExtractBSPvalue* function, which assigns *p*-values to each CpG based on binomial tests incorporating the raw read counts and the sample-specific bisulfite conversion rate and uses the Benjamini-Hochberg step-up procedure to control the false discovery rate for multiple testing. We specified an accepted error interval of 0.2 (the default value) and an FDR of 0.05. Only significant sites were used in downstream analyses. For specimens with fewer than 200 phage cytosines analyzed (5 of the 75 specimens) we used the minimum bisulfite conversion rate from other specimens from the same ddRAD sublibrary, which were pooled together in the same bisulfite conversion reaction and should have the same conversion rate.

### Data analysis

To assess data yield in specimens of different ages, we modeled the total number of cleaned reads (demultiplexed and trimmed) and aligned reads per specimen. We also modeled the number of unique CpG positions sequenced per specimen. These analyses were done using negative binomial regression implemented in *R* v.3.5.1 (RCoreTeam 2018) with the *glm.nb* function of the package *mass* v.7.3-50 (Ripley et al. 2013). We modeled each measure separately with fixed effects of species and specimen age. We used Tukey tests for all pairwise comparisons, implemented using the *glht* function of the *R* package *multcomp* v.1.4-8 (Hothorn et al. 2014). We report Bonferroni corrected *p*-values for all pairwise comparisons.

To characterize percent methylation, we modeled raw read counts of methylated and unmethylated cytosines at each locus using binomial generalized linear mixed models with a logit link function and fit with Laplace approximation, implemented using the *glmer* function of the *R* package *lme4* v.1.1-20 (Bates et al. 2014). Because cytosine methylation shows high spatial correlation (Eckhardt et al. 2006), data from CpGs occurring within 1000 bp of each other in the genome were pooled into a single locus. Sequences with a read depth less than 10X were excluded following conservative guidelines for calling percent methylation (Ziller et al. 2015). To account for PCR duplication, we also excluded positions with abnormally high coverage, defined as bases in the top 99.9th percentile of read depth for each individual (following Hu et al. 2018). Because many loci were sequenced for each individual, we included specimen identity as a random intercept term in all models. We also included an observation-level random effect in all models to account for overdispersion (Harrison 2014). Dispersion parameters are reported for each model below.

To test for abnormalities in methylation calling associated with specimen age, we used Mbias plots to check for biased methylation estimates toward read ends and compared global methylation estimates due to specimen age. To produce the Mbias plots, Bismark M-bias report files for each specimen were combined and visualized using the *MethylationTuples* v.0.3.0 package in *R* (Hickey 2015). To assess global methylation, we modeled methylation at each locus with species and specimen age as fixed effects (dispersion parameter = 1.020). For this analysis, all loci including known autosomal loci, known X chromosome loci, and unplaced loci were included; because the reference scaffold for *P. leucopus* lacks chromosome assignments, sex chromosomes could not be omitted from the analysis.

Finally, we tested whether methylation estimates followed predicted patterns for mammalian methylation in known genomic regions; namely, we compared putative promoters versus gene bodies and autosomes versus X chromosomes. These analyses were only done for *P. maniculatus* because the reference genome for *P. leucopus* lacks annotations and chromosome assignments. We first compared methylation estimates in promoters, which we predicted would show reduced methylation, and gene bodies, which we predicted would show increased methylation. We modeled methylation with genomic region and specimen age as fixed effects and compared methylation in promoters and gene bodies (dispersion parameter = 1.041). Genomic regions were defined by sequence annotations downloaded from *Ensembl* (the pbairdii_gene_ensembl dataset) following the classification method outlined in Pedersen et al. (2014). Briefly, putative promoter regions were defined as the region 500 bp upstream and 2000 bp downstream of the transcription start site (TSS) for the first exon in a gene, gene bodies were defined as the region from the end of the promoter (2000 bp downstream from the TSS) to the final transcription end position in the gene, and all loci not defined as promoters or gene bodies were labelled as other. *Ensembl* annotations were downloaded and processed using the *R* package *biomaRt* v.2.36.1 (Kinsella et al. 2011). To assess chromosome methylation, we compared locus methylation in autosomes, female X chromosomes, and male X chromosomes. This analysis was only performed for *P. maniculatus* from the youngest age group (0-3 yo) because older specimens did not yield enough loci from the X chromosome. We modeled methylation at each locus with chromosome type as a fixed effect (dispersion parameter = 1.042).

## Results

### Bisulfite conversion efficiency

The bisulfite conversion rates calculated from the lambda phage reads indicated almost complete conversion in all samples (sequencing statistics for each specimen are included in Supplementary Table S1). The 0.1% phage spike-in produced a sufficient number of cytosines (over 200) to estimate conversion efficiency in all but five (out of 75) samples; for those samples, the average conversion rate of the sublibrary was used for methylation calling as described in *Methods - Methylation calling*. After adjusting for low coverage, estimated conversion rates ranged from 94.2% - 100% (mean 98.9%).

### Data yield

For all three measures of data yield, younger specimens yielded more data than older specimens (Table 1). The total number of cleaned read pairs, defined as pairs retained after demultiplexing and trimming for quality, was greater in 0-3 yo specimens than 13 yo specimens (1.379±0.334; *z* = 4.132, *p* = 0.0001) and 76 yo specimens (2.352±0.359; *z* = 6.555, *p <* 0.0001) and was greater in 13 yo specimens than 76 yo specimens (0.973±0.358; *z* = 2.718, *p* = 0.020). The number of cleaned 0.973 0.358 read pairs did not differ between the two species (*z* = −1.259, *p* = 0.208). The total number of aligned read pairs, defined as pairs retained after aligning to the reference genome, also increased with each time step; more aligned pairs were retained for 0-3 yo specimens than 13 yo specimens (2.229±0.405; *z* = 5.509, *p <* 0.0001) and 76 yo specimens (3.295±0.435; *z* = 7.574, *p <* 0.0001) and more pairs were retained for 13 yo specimens than 76 yo specimens (1.066±0.434; *z* = 2.457, *p* = 0.042). Significantly more aligned read pairs were retained for *Peromyscus leucopus* specimens than *Peromyscus maniculatus* specimens (0.846±0.346; *z* = 2.444, *p <* 0.015). The total number of CpG positions sequenced was greater in 0-3 yo specimens than 13 yo specimens (2.652±0.352; *z* = 7.534, *p <* 0.0001) and 76 yo specimens (3.923±0.378; *z* = 10.368, *p <* 0.0001) and was greater in 13 yo specimens than 76 yo specimens (3.923±0.378; *z* = 3.369, *p* = 0.002) (Figure 2).

**Table 1:** Sequencing statistics grouped by species and year collected. The number of specimens is indicated in the N spec column. The total number of read pairs sequenced is shown for cleaned reads (pairs retained after demultiplexing and cleaning) and aligned reads (pairs retained after alignment to the reference genome). The total number of CpG positions is shown for a minimum read depth of 1X, 5X, and 10X.

| Species | Age | Year | N spec | Read pairs |  | CpG positions |  |  |
| --- | --- | --- | --- | --- | --- | --- | --- | --- |
|  |  |  |  | Cleaned | Aligned | 1X | 5X | 10X |
| *P. leucopus* | 76 | 1940 | 13 | 1053049 | 292168 | 26714 | 6185 | 5463 |
|  | 13 | 2003 | 14 | 3640337 | 958386 | 125380 | 32102 | 28769 |
|  | 1-3 | 2013-15 | 13 | 4690267 | 2231013 | 498326 | 166270 | 75872 |
| *P. maniculatus* | 76 | 1940 | 8 | 318805 | 33443 | 5761 | 1494 | 1312 |
|  | 13 | 2003 | 13 | 1042870 | 181536 | 24871 | 1388 | 1154 |
|  | 0 | 2016 | 14 | 10961871 | 4552427 | 1025598 | 518267 | 277450 |

**Figure 2:**
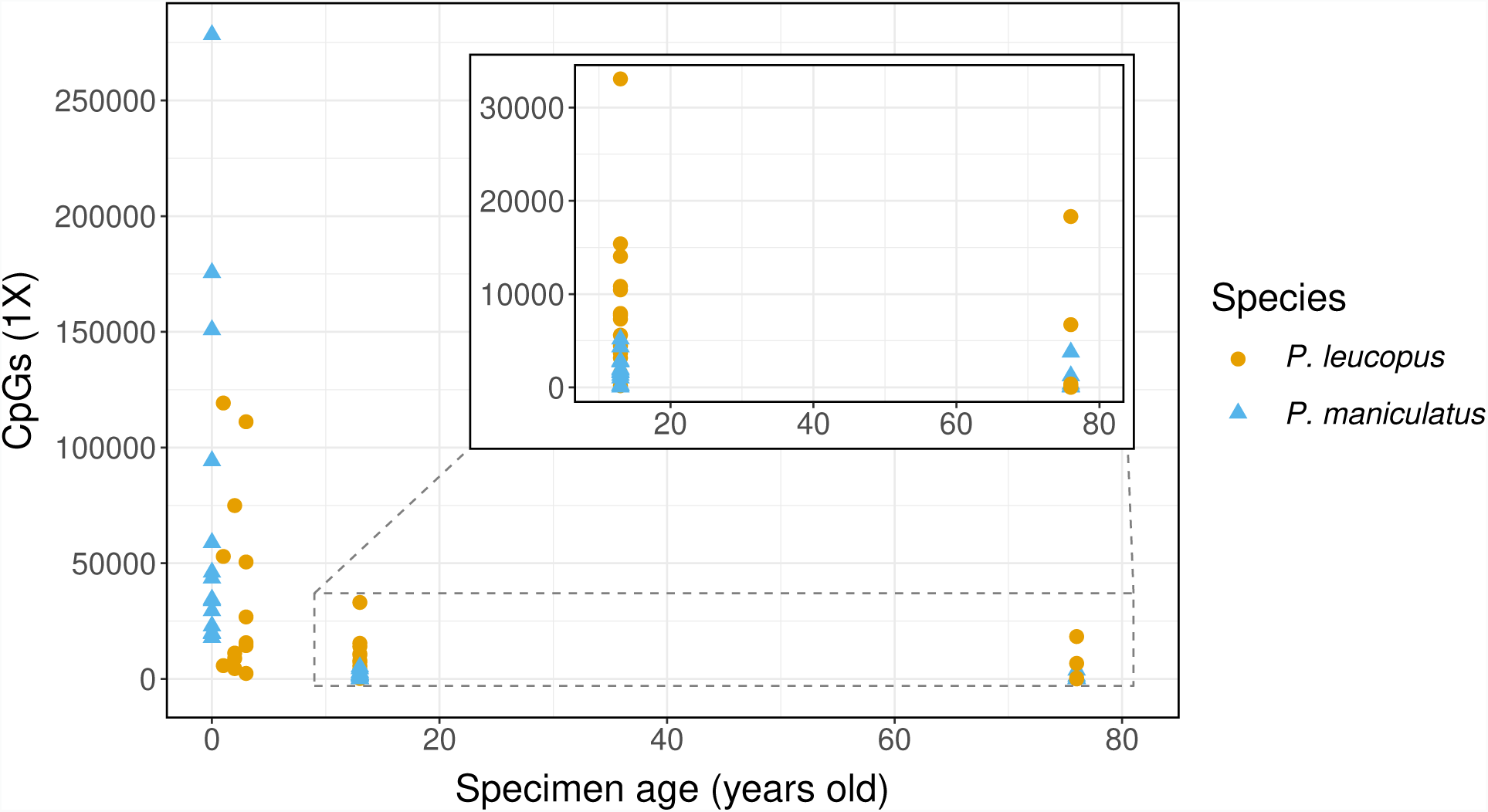
Total CpGs sequenced per specimen by specimen age (minimum depth = 1X). Orange circles indicate *P. leucopus* and blue triangles indicate *P. maniculatus*. Inset: Zoomed view of specimens 13 - 76 years old (area shown in the dashed box).

### Global methylation estimates

Plots of percent methylation at each position along reads were visually assessed for read end biases (Supplementary Figure S2). Reads were trimmed for cut site sequences (first 5 positions of forward reads and first 4 positions of reverse reads) and forward reads were further trimmed for quality by removing 5 bp at the 5’ end and truncating reads at 118 bp. After trimming, these plots revealed greater variation in older versus newer specimens but no systematic methylation biases due to read position.

Estimated global methylation rates were significantly lower in *P. maniculatus* than in *P. leucopus*(−0.518±0.085; *z* = −6.076, *p* < 0.0001; odds ratio (OR) = 0.596); average methylation over all loci was 64.3% and 67.1%, respectively. Global methylation estimates were significantly lower in the oldest age group (76 yo) than in the youngest age group (0-3 yo) (−291±0.119; *z* = −2:437, *p* = 0.044; OR = 0.748). No significant differences in methylation estimates were observed between 13 yo specimens and 76 yo specimens (*z* = 0.706, *p* = 0.480) or 1-3 yo specimens (*z* = 1.461, *p* = 0.144). In both species in all age groups, locus methylation followed a bimodal distribution in which fully methylated (100%) and fully unmethylated (0%) loci were more common than partially methylated loci (Figure 3).

**Figure 3:**
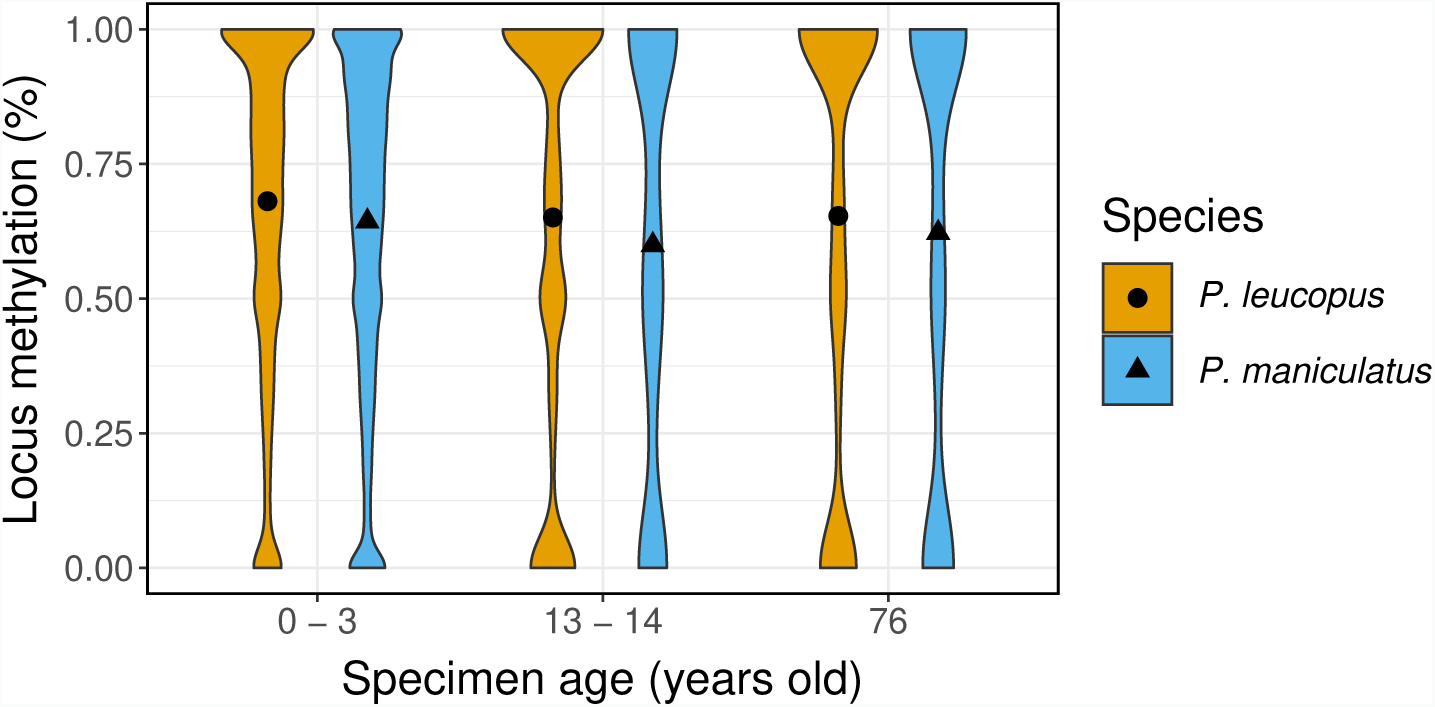
Violin plot of percent methylation across all loci by specimen age and species. Global methylation rates were signficantly reduced in *P. maniculatus* relative to *P. leucopus*. Methylation rates were also reduced in 76 year old specimens relative to 0-3 year old specimens.

### Methylation in known genomic regions in *P. maniculatus*

Methylation rates varied between different genomic regions following expected trends for mammalian genomes. Methylation was greater in gene bodies relative to promoter regions (1.297±0.039; *z* = 33.38, *p <* 0.0001; OR = 3.658; Fig 4). Average locus methylation was 51.4% in promoter regions and 68.2% in gene body regions. Regional methylation did not differ significantly due to specimen age (relative to 0-3 yo specimens, 13 yo specimens: *z* = 0.619, *p* = 0.536; 76 yo: *z* = 0.998, *p* = 0.318).

**Figure 4:**
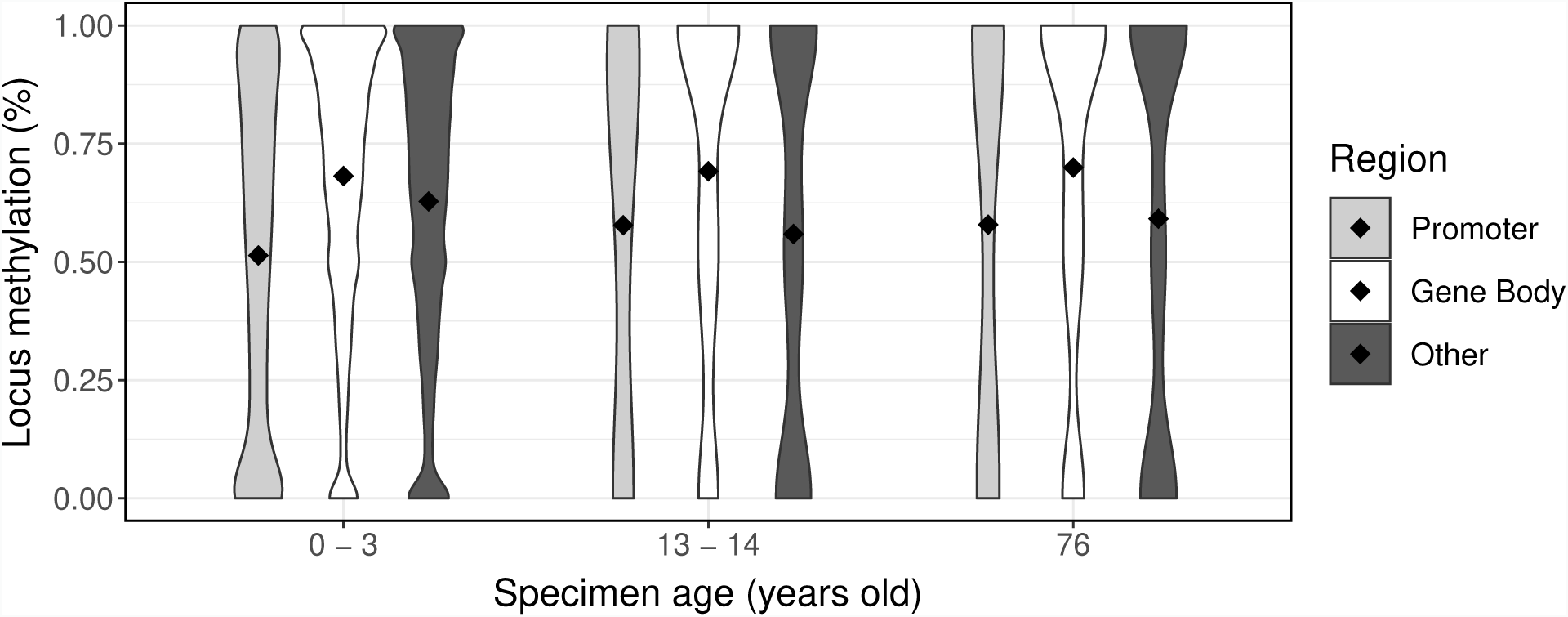
Violin plot of percent methylation in putative promoters, gene bodies, and unknown genomic regions (Other) in each age group. Methylation in promoters was reduced relative to methylation in gene bodies.

Chromosome-specific patterns could only be assessed in *P. maniculatus* from the youngest age group; older specimens did not yield enough loci from the X chromosome to describe the distribution of locus methylation. Loci from autosomes and the male X chromosome followed a bimodal distribution in percent methylation; fully methylated (100%) and fully unmethylated (0%) loci were more common than partially methylated loci. Loci from the female X chromosome showed reduced bimodality, with fewer fully methylated and fully unmethylated loci and more loci with intermediate methylation (Figure 5). Average locus methylation was reduced in female X chromosomes relative to autosomes (*−*0.681±0.082; *z* = *−*8.296, *p <* 0.0001; odds ratio (OR) = 0.506) and was increased in male X chromosomes relative to autosomes (0.271±0.066 *z* = 4.128 *p <* 0.0001 odds ratio (OR) = 1.311). Average methylation over all loci was 64.6% for autosomes, 54.8% for female X chromosomes, and 67.3% for male X chromosomes.

**Figure 5:**
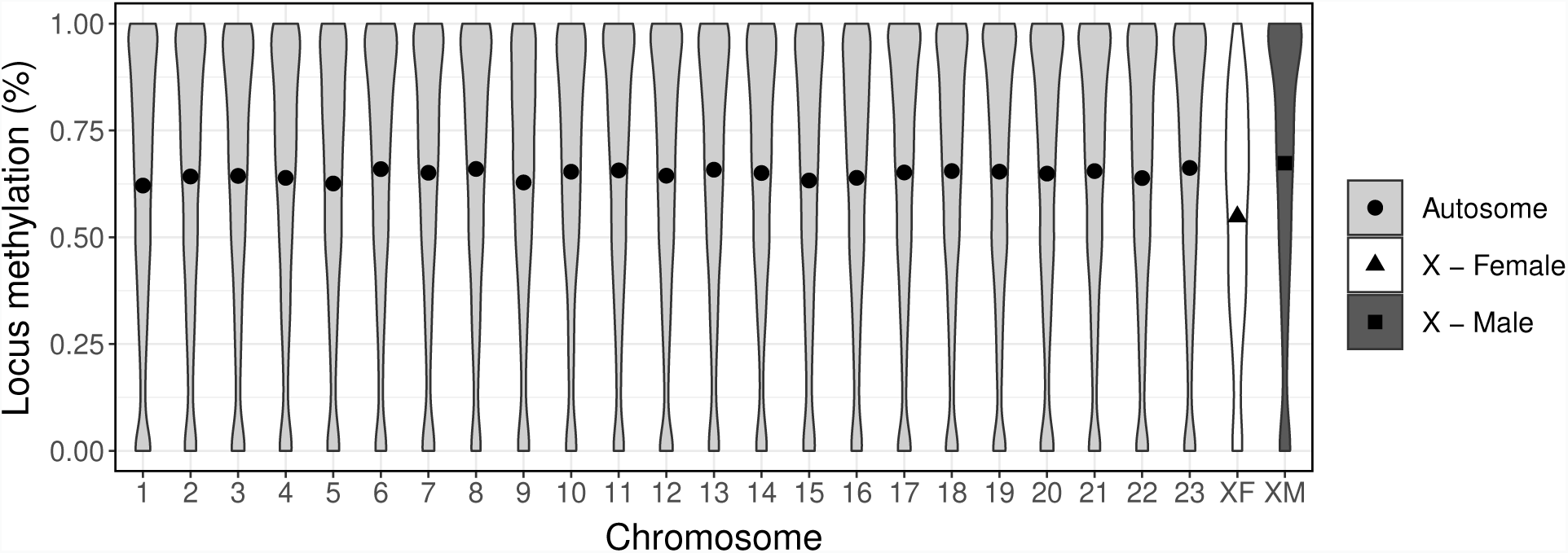
Distribution of locus methylation in all autosomes, female X chromosomes, and male X chromosomes in *P. maniculatus* collected in 2016. Relative to autosomes, methylation was significantly reduced in the female X chromosome and significantly increased in the male X chromosome.

## Discussion

The cytosine methylation patterns we recovered from dried skull specimens, including samples up to 76 years old, demonstrate the enormous resource contained in natural history collections. However, our dataset also highlights the challenges of conducting epigenetic studies using historic samples. As in museum genomic studies, museum epigenomic studies must account for reduced yield and high variability in the data produced by historic specimens. These issues are discussed in more detail below.

### Variability in specimen yield

Older specimens yielded less data than younger specimens, and data yield is likely to be the primary challenge for future studies that use historic museum specimens. However, our results indicate that some older specimens perform well; for example, our two 76 yo specimens with the highest extracted DNA concentrations (over 9 ng/*µ*L) sequenced a number of CpG positions comparable to specimens in the 13-14 yo and 0-3 yo age groups (Figure 2). This disparity in specimen performance is typical of older historic specimens, which tend to show high variation in the quantity of recoverable DNA.

Our results suggest that starting DNA concentration may be a better predictor of performance than specimen age. In addition, both of our high quality 76 yo samples were diluted to standardize concentration during library preparation, suggesting that they could potentially yield more CpGs if prepared at a higher concentration. Other options for increasing data yield are discussed below.

### Global methylation estimates

To test for abnormalities in methylation calling in older specimens, we assessed our data for methylation biases near read ends and modeled global methylation levels as a function of specimen age. In particular, we tested for a signal of *post mortem* hydrolytic deamination, which causes the spontaneous conversion of cytosine into either uracil (in the case of unmethylated cytosine) or thymine (in the case of 5-methylcytosine) (Willerslev and Cooper 2005). In ancient or historic genomics studies, this conversion results in erroneous C to T SNP calling; in bisulfite studies, deaminated cytosines could be misinterpreted as unmethylated cytosines and cause depressed methylation estimates for older specimens. Deamination tends occur at higher rates near read ends, however, we did not observe such a signal in our reads in any age group (Supplementary Figure S2). The lack of read end deamination was likely an outcome of sampling the genome using double digestion. Deamination tends to occur near the ends of fragmented DNA where single strand overhangs occur, however, these natural breaks are less likely to be sequenced when two restriction enzymes are used to cleave the DNA at each end. The methylation bias plots also revealed more variation in methylation estimates at each read position in older specimens. This variation probably reflects the lower number of reads averaged at each position for older specimens rather than systematic biases within the dataset.

Our estimates of global methylation may indicate an effect of deamination in our oldest age group. Methylation in 76 yo specimens was reduced relative to 0-3 yo specimens, though the effect was marginally significant (p=0.044). The odds ratio of 0.748 indicated that the likelihood of calling a given CpG position as methylated is about 25% less likely in 76 yo specimens relative to 0-3 yo specimens. Assuming that the true methylation level does not vary between the mice sampled in 1940 and 2013-2016, our results suggest that deamination may bias methylation estimates in older historic specimens even in protocols such as ours with minimal read end deamination. Future studies should test for a potential signal of deamination and take steps to reduce sequencing of deaminated sites. For example, uracil-DNA-glycosylase and endonuclease VIII can be used to remove uracil prior to bisulfite treatment, which will avoid miscalled bases due to deaminated unmethylated cytosines (though not methylated cytosines) (Briggs et al. 2010).

### Methylation of known genomic regions in *P. maniculatus*

The observed patterns in known genomic regions were consistent with expectations for *in vivo* methylation of mammalian somatic cells. A CpG dinucleotide within a gene body was over 3.5 times as likely to be methylated as a CpG within a putative promoter region (odds ratio = 3.658). This pattern of reduced methylation in promoters and increased methylation in coding regions is consistent with expectations for mammalian DNA (Jones 2012). Locus methylation in autosomes showed a bimodal distribution with peaks at 0% and 100%, as is expected for autosomal loci within a single cell type (Rakyan et al. 2004; Eckhardt et al. 2006). Loci in the male X chromosome showed a similar bimodal distribution, but loci in the female X chromosome showed a decreased frequency of fully methylated and fully unmethylated loci and an increased frequency of loci with intermediate methylation. Duncan et al. (2018) described similar methylation distributions across autosomes, female X chromosomes, and male X chromosomes in liver cells of *Mus musculus*. The reduced bimodality observed in female X chromosomes likely reflects the role of methylation in X-inactivation, a mechanism of dosage compensation in female mammals. Loci that undergo X-inactivation are often hypermethylated on the inactive X and hypomethylated on the active X, resulting in intermediate measures of methylation when data from the two chromosomes are aggregated (Hellman 2007).

### Increasing the success of epigenomic studies based on historic samples

Probably the greatest challenge to museum epigenomics studies will be reduced sequencing success in historic specimens due to low DNA concentration or DNA fragmentation. Several steps of our bisulfite ddRAD protocol could be modified or replaced to increase yield from historic specimens. For example, the size selection window could be reduced to compensate for fragmentation in historic DNA. Selecting for smaller fragments may increase yield, though the gain in loci will be accompanied by reduced numbers of homologous loci sequenced across individuals. Steps could also be taken to minimize DNA degradation during the bisulfite treatment; for example, shortening the bisulfite incubation time should reduce DNA damage, though it may also reduce conversion efficiency (Grunau 2002). Our protocol also cleaved the DNA with two restriction enzymes, which may have contributed to problems in amplification and sequencing associated with DNA fragmentation. However, double digestion may also minimize the signal of read-end deamination, as discussed above.

Many genomic library preparation protocols have been described for increasing yield from damaged and fragmented DNA. For example, libraries can be prepared without digestion or sonication and sequenced directly to avoid further fragmentation (Burrell et al. 2015), or low input bisulfite methods can be used when limited DNA is available (Miura and Ito 2018). Enrichment methods seem to be particularly effective for sampling degraded historic and ancient DNA (Jones and Good 2016; Suchan et al. 2016). Seguin-Orlando et al. (2015) described methylation-based enrichment methods for ancient DNA which may be promising for museum epigenomic work, though the authors outline biases in template enrichment that should be considered (*e.g.*, longer fragments and regions with limited deamination). Methylation-based enrichment also selectively targets CpG-rich regions, as does traditional reduced representation bisulfite sequencing; such protocols may be more fitting for studies focusing on regulatory regions such as CpG islands and promoters. Alternatively, it may be possible to avoid bisulfite conversion altogether; several ancient epigenomics studies have reconstructed methylation maps from patterns of hydrolytic deamination (*e.g.*, Briggs et al. 2010; Gokhman et al. 2014; Pedersen et al. 2014; Hanghøj et al. 2016). This approach would not have been possible for our specimens, which did not show a strong deamination signal, however it may be an option for museum specimens that are highly degraded. In addition, several cheaper options are available for measuring methylation at fewer sites, such as MS-AFLP and targeted bisulfite sequencing; for example, Smith et al. (2015) used targeted bisulfite pyrosequencing to describe methylation at an imprinted site in ancient humans.

Museum epigenomics studies will need to accommodate the large variance in the quantity of data produced by individual historic specimens. Sampling designs should account for a high failure rate in older specimens, or if possible, specimens should be screened in advance of library preparation for DNA quantity and quality (for example, by characterizing fragment size distributions). We expect that most samples that can be used for genomic work can also be used for epigenomic work. Because high quality specimens are likely to be rare, analyses that require fewer individuals will probably be more successful.

### Applications of epigenomic data from historic specimens

Methylation is one of the best-studied epigenetic mechanisms and is associated with a range of processes, from development to disease response to phenotypic plasticity. One of the most intriguing directions for museum epigenomics research is the study of characteristics that do not fossilize, such as non-morphological traits or historical environmental conditions. For example, methylation variation modulates gene expression related to various behavioral (*e.g.*, Meaney and Szyf 2005) and physiological traits (*e.g.*, García-Carpizo et al. 2011). Murphy and Benítez-Burraco (2018) used methylation patterns to infer the expression of language processing genes in Neanderthals. Environmentally-induced methylation variation can reflect environmental conditions such as food availability (*e.g.*, Heijmans et al. 2008), climate (*e.g.*, Fu et al. 2010; Gugger et al. 2016), and exposure to disease or toxins (Robertson 2005; Baccarelli and Bollati 2009). Gokhman et al. (2017) demonstrated how methylation patterns can be used to study past environments by describing markers of prenatal nutrition in Denisovan and Neanderthal genomes. Ancient and historic epigenomic studies may allow us to explore aspects of past populations that are not reflected in a specimen’s morphology or genetic sequence.

Museum epigenomics studies also provide the opportunity to directly measure how epigenetic effects change over time. Just as in museum genomic studies (Burrell et al. 2015), epigenomic studies can use collections to describe temporal changes in population-level variation. Such studies could help clarify a range of unresolved questions in ecological epigenetics, including the transgenerational stability of epigenetic marks, the timescales of induction of epigenetic effects, and the relationship between epigenetic and genetic variation. It is still unclear what role, if any, non-genetic mechanisms such as epigenetic effects play in evolutionary processes (*e.g.*, Laland et al. 2014). Observing change over time in epigenetic effects may provide insights into their role in adaptation and evolution.

## Acknowledgements

We thank the Dantzer lab and Knowles lab members for their support and help, especially Freya van Kesteren and Andrea Thomaz. We thank Susan Hoffman and Joe Baumgartner at Miami University for sharing specimens and expertise, and students at University of Michigan (Austin Rife, Daniel Nondorf, Anne Sabol, Francesca Santicchia) and Miami University (Jeremy Papuga) for assisting with specimen collection. We thank the University of Michigan Museum of Zoology for providing specimens and the Genomic Diversity Lab at the University of Michigan for hosting the genomic work in their ancient DNA facility. We thank Emiliano Trucchi for advice about the protocol and methylated adapter design. We thank Phil Myers, Cody Thompson, and Raquel Rivadeneira for their support. We also thank three anonymous reviewers for their helpful comments. TLR was funded by the National Science Foundation Postdoctoral Research Fellowship in Biology (Award #: 1612143). This work was funded in part by the University of Michigan (BD, LLK) and the NSF PRFB.

## Author Contributions

Conceived the experiment: TLR and BD. Designed the experiment: TLR, BD, and LLK. Contributed reagents: TLR, BD, and LLK. Performed the research and data analysis: TLR. Wrote the manuscript: TLR, BD, and LLK.

