## Supplemental Figures for "Museum epigenomics: characterizing cytosine methylation in historic museum specimens"

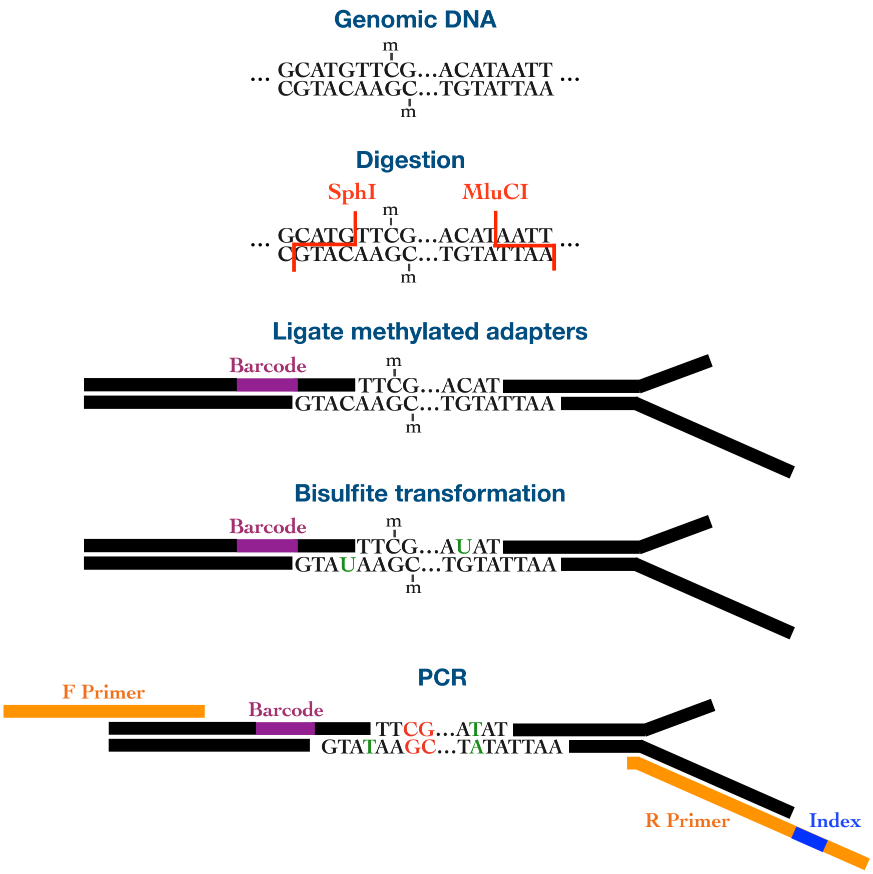


**Figure S1:** Flowchart of bisulfite ddRAD library preparation protocol. **1. Genomic DNA.** Genomic DNA with methylated cytosines indicated with -m. **2. Digestion.** DNA is digested with restriction enzymes SphI-HF and MluCI. **3. Ligate methylated adapters.** Methylated barcoded adapters are ligated onto the digested DNA fragments. **4. Bisulfite transformation.** Bisulfite treatment transforms unmethylated cytosines to uracil. Methylated cytosines remain unchanged. **5. PCR.** DNA is amplified using indexed primers. Uracils are replaced by thymines. Red bases indicate methylated cytosines or their complements, and green bases indicate unmethylated cytosines (now thymines) or their complements.

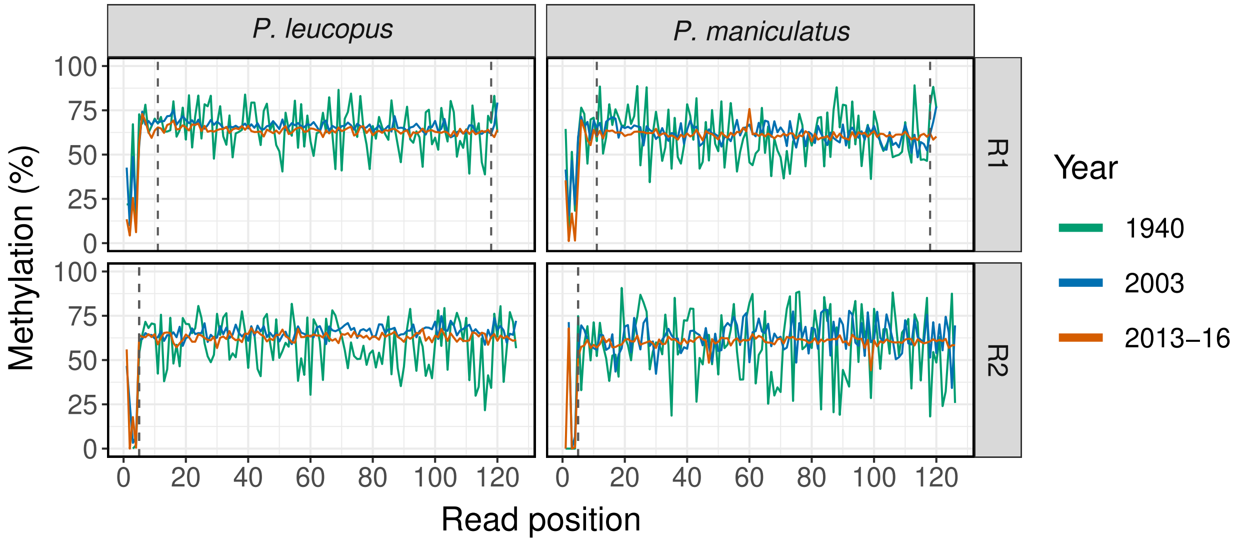


**Figure S2:** MBias plot of percent methylation at each read position for both species in all three year groups. The first 5 positions on the 5' end of forward reads (top panels) and the first 4 positions on the 5' end of reverse reads (bottom panels) correspond to the cut site sequences. These bases were trimmed before aligning reads to the genome. Forward reads were further trimmed for read-end quality by removing 5 bp from the 5' end and truncating reads at 118 bp at the 3' end (cutoffs are indicated by the dashed lines).
