## Supplemental Tables for "Museum epigenomics: characterizing cytosine methylation in historic museum specimens"

**Table S1:** Oligonucleotides used in this protocol. All cytosines in RAD adapters are methylated (indicated by C$^m$). Phos indicates a phosphate group.

| Type | Name | Sequence |
|---|---|---|
| Barcoded methylated RAD adapters | GCATG_P1.1m | AC$^m$AC$^m$TC$^m$TTTC$^m$C$^m$C$^m$TAC$^m$AC$^m$GAC$^m$GC$^m$TC$^m$TTC$^m$C$^m$GATC$^m$TGC$^m$ATGC$^m$ATG |
| | AACCA_P1.1m | AC$^m$AC$^m$TC$^m$TTTC$^m$C$^m$C$^m$TAC$^m$AC$^m$GAC$^m$GC$^m$TC$^m$TTC$^m$C$^m$GATC$^m$TAAC$^m$C$^m$AC$^m$ATG |
| | CGATC_P1.1m | AC$^m$AC$^m$TC$^m$TTTC$^m$C$^m$C$^m$TAC$^m$AC$^m$GAC$^m$GC$^m$TC$^m$TTC$^m$C$^m$GATC$^m$TC$^m$GATC$^m$C$^m$ATG |
| | TCGAT_P1.1m | AC$^m$AC$^m$TC$^m$TTTC$^m$C$^m$C$^m$TAC$^m$AC$^m$GAC$^m$GC$^m$TC$^m$TTC$^m$C$^m$GATC$^m$TTC$^m$GATC$^m$ATG |
| | TGCAT_P1.1m | AC$^m$AC$^m$TC$^m$TTTC$^m$C$^m$C$^m$TAC$^m$AC$^m$GAC$^m$GC$^m$TC$^m$TTC$^m$C$^m$GATC$^m$TTGC$^m$ATC$^m$ATG |
| | CAACC_P1.1m | AC$^m$AC$^m$TC$^m$TTTC$^m$C$^m$C$^m$TAC$^m$AC$^m$GAC$^m$GC$^m$TC$^m$TTC$^m$C$^m$GATC$^m$TC$^m$AAC$^m$C$^m$C$^m$ATG |
| | GGTTG_P1.1m | AC$^m$AC$^m$TC$^m$TTTC$^m$C$^m$C$^m$TAC$^m$AC$^m$GAC$^m$GC$^m$TC$^m$TTC$^m$C$^m$GATC$^m$TGGTTGC$^m$ATG |
| | AAGGA_P1.1m | AC$^m$AC$^m$TC$^m$TTTC$^m$C$^m$C$^m$TAC$^m$AC$^m$GAC$^m$GC$^m$TC$^m$TTC$^m$C$^m$GATC$^m$TAAGGAC$^m$ATG |
| | AGCTA_P1.1m | AC$^m$AC$^m$TC$^m$TTTC$^m$C$^m$C$^m$TAC$^m$AC$^m$GAC$^m$GC$^m$TC$^m$TTC$^m$C$^m$GATC$^m$TAGC$^m$TAC$^m$ATG |
| | ACACA_P1.1m | AC$^m$AC$^m$TC$^m$TTTC$^m$C$^m$C$^m$TAC$^m$AC$^m$GAC$^m$GC$^m$TC$^m$TTC$^m$C$^m$GATC$^m$TAC$^m$AC$^m$AC$^m$ATG |
| | GCATG_P1.2m | [Phos]C$^m$ATGC$^m$AGATC$^m$GGAAGAGC$^m$GTC$^m$GTGTAGGGAAAGAGTGT |
| | AACCA_P1.2m | [Phos]TGGTTAGATC$^m$GGAAGAGC$^m$GTC$^m$GTGTAGGGAAAGAGTGT |
| | CGATC_P1.2m | [Phos]GATC$^m$GAGATC$^m$GGAAGAGC$^m$GTC$^m$GTGTAGGGAAAGAGTGT |
| | TCGAT_P1.2m | [Phos]ATC$^m$GAAGATC$^m$GGAAGAGC$^m$GTC$^m$GTGTAGGGAAAGAGTGT |
| | TGCAT_P1.2m | [Phos]ATGC$^m$AAGATC$^m$GGAAGAGC$^m$GTC$^m$GTGTAGGGAAAGAGTGT |
| | CAACC_P1.2m | [Phos]GGTTGAGATC$^m$GGAAGAGC$^m$GTC$^m$GTGTAGGGAAAGAGTGT |
| | GGTTG_P1.2m | [Phos]C$^m$AAC$^m$C$^m$AGATC$^m$GGAAGAGC$^m$GTC$^m$GTGTAGGGAAAGAGTGT |
| | AAGGA_P1.2m | [Phos]TC$^m$C$^m$TTAGATC$^m$GGAAGAGC$^m$GTC$^m$GTGTAGGGAAAGAGTGT |
| | AGCTA_P1.2m | [Phos]TAGC$^m$TAGATC$^m$GGAAGAGC$^m$GTC$^m$GTGTAGGGAAAGAGTGT |
| | ACACA_P1.2m | [Phos]TGTGTAGATC$^m$GGAAGAGC$^m$GTC$^m$GTGTAGGGAAAGAGTGT |
| | flex_P2.1m | GTGAC$^m$TGGAGTTC$^m$AGAC$^m$GTGTGC$^m$TC$^m$TTC$^m$C$^m$GATC$^m$T |
| | flex_P2.2m | [Phos]AATTAGATC$^m$GGAAGAGC$^m$GAGAAC$^m$AA |
| Indexed PCR primers | PCR1 | AATGATACGGCGACCACCGAGATCTACACTCTTTCCCTACACGACG |
| | PCR2_ATCACG | CAAGCAGAAGACGGCATACGAGATCGTGATGTGACTGGAGTTCAGACGTGTGC |
| | PCR2_CGATGT | CAAGCAGAAGACGGCATACGAGATACATCGGTGACTGGAGTTCAGACGTGTGC |
| | PCR2_TTAGGC | CAAGCAGAAGACGGCATACGAGATGCCTAAGTGACTGGAGTTCAGACGTGTGC |
| | PCR2_TGACCA | CAAGCAGAAGACGGCATACGAGATTGGTCAGTGACTGGAGTTCAGACGTGTGC |
| | PCR2_ACAGTG | CAAGCAGAAGACGGCATACGAGATCACTGTGTGACTGGAGTTCAGACGTGTGC |
| | PCR2_GCCAAT | CAAGCAGAAGACGGCATACGAGATATTGGCGTGACTGGAGTTCAGACGTGTGC |
| | PCR2_CAGATC | CAAGCAGAAGACGGCATACGAGATGATCTGGTGACTGGAGTTCAGACGTGTGC |
| | PCR2_ACTTGA | CAAGCAGAAGACGGCATACGAGATTCAAGTGTGACTGGAGTTCAGACGTGTGC |
| | PCR2_GATCAG | CAAGCAGAAGACGGCATACGAGATCTGATCGTGACTGGAGTTCAGACGTGTGC |
| | PCR2_TAGCTT | CAAGCAGAAGACGGCATACGAGATAAGCTAGTGACTGGAGTTCAGACGTGTGC |

**Table S2:** Specimen information and sequencing statistics for all specimens used in this study. Specimens beginning with MZ were sampled from the University of Michigan Museum of Zoology collection. The Age column indicates the specimen age at the time of DNA extraction (years old). Sequencing statistics indicate the concentration of DNA after extraction (Ext DNA conc), the starting amount of DNA used for library preparation (Starting DNA amt), the sample-specific bisulfite conversion rate measured using a spike-in of fully unmethylated phage DNA (Bisulfite conv rate), the number of read pairs retained after demultiplexing and cleaning (Demult read pairs), the number of read pairs retained after alignment (Aligned read pairs), the mapping efficiency (Mapping eff), and the total number of unique CpG positions sequenced at a read depth of 1X and 10X. The starred specimens were sequenced in two separate libraries and the reads from the two libraries were combined prior to methylation calling; therefore, the number of CpG positions reported in the last two columns represent this combined value.

| Specimen | Species | Sex | Date collected | Age (yo) | County | Location | Ext DNA conc (ng/$\mu$L) | Starting DNA amt (ng) | Bisulfite conv rate | Demult read pairs | Aligned read pairs | Mapping eff (%) | CpG pos (1X) | CpG pos (10X) |
|---|---|---|---|---|---|---|---|---|---|---|---|---|---|---|
| 13MN007 | Pln | M | 8/6/13 | 3 | Menominee | 45.52,-87.41 | 33.90 | 350 | 0.99 | 1205740 | 606793 | 50.30 | 111150 | 14258 |
| 13MN031 | Pln | M | 8/7/13 | 3 | Menominee | 45.48,-87.42 | 14.40 | 350 | 1.00 | 59625 | 22876 | 38.40 | 15672 | NA |
| 13MN033 | Pln | F | 8/7/13 | 3 | Menominee | 45.48,-87.42 | 15.40 | 350 | 1.00 | 56337 | 20381 | 36.20 | 14416 | NA |
| 13MN035 | Pln | M | 8/7/13 | 3 | Menominee | 45.48,-87.42 | 35.00 | 350 | 0.99 | 189141 | 100893 | 53.30 | 26793 | 529 |
| 13MN040 | Pln | F | 8/8/13 | 3 | Menominee | 45.48,-87.42 | 34.20 | 350 | 0.99 | 716689 | 395173 | 55.10 | 50551 | 18155 |
| 13MN045 | Pln | F | 8/8/13 | 3 | Menominee | 45.48,-87.42 | 23.60 | 350 | 0.99 | 25412 | 7627 | 30.00 | 2381 | 39 |
| 14MN003 | Pln | M | 8/12/14 | 2 | Menominee | 45.52,-87.41 | 26.80 | 350 | 1.00 | 35995 | 15056 | 41.80 | 8884 | 33 |
| 14MN005 | Pln | F | 8/12/14 | 2 | Menominee | 45.48,-87.42 | 28.10 | 350 | 1.00 | 512745 | 247837 | 48.30 | 74927 | 6087 |
| 14MN010 | Pln | M | 8/12/14 | 2 | Menominee | 45.48,-87.42 | 4.14 | 350 | 1.00 | 58085 | 19408 | 33.40 | 11171 | 8 |
| 14MN024 | Pln | F | 8/13/14 | 2 | Menominee | 45.48,-87.42 | 34.50 | 350 | 0.99 | 29078 | 10629 | 36.60 | 4464 | 12 |
| 15MN001 | Pln | F | 7/24/15 | 1 | Menominee | 45.48,-87.42 | 10.60 | 350 | 0.99 | 63300 | 20337 | 32.10 | 5754 | 258 |
| 15MN002 | Pln | M | 7/24/15 | 1 | Menominee | 45.48,-87.42 | 49.20 | 350 | 0.99 | 601948 | 211803 | 35.20 | 52939 | 8451 |
| 15MN003 | Pln | M | 7/24/15 | 1 | Menominee | 45.48,-87.42 | 18.90 | 350 | 1.00 | 1136172 | 552200 | 48.60 | 119224 | 28042 |
| MZ11320 | Pmg | M | 10/22/16 | 0 | Menominee | 45.5,-87.41 | 350.00 | 350 | 1.00 | 3252521 | 1467424 | 45.10 | 278344 | 91530 |
| MZ11321 | Pmg | M | 10/22/16 | 0 | Menominee | 45.5,-87.41 | 54.00 | 350 | 1.00 | 167673 | 73602 | 43.90 | 34418 | 204 |
| MZ11322 | Pmg | F | 10/22/16 | 0 | Menominee | 45.5,-87.41 | 40.30 | 350 | 1.00 | 221935 | 97250 | 43.80 | 43589 | 867 |
| MZ11323 | Pmg | M | 10/22/16 | 0 | Menominee | 45.5,-87.41 | 41.50 | 350 | 1.00 | 338886 | 149205 | 44.00 | 58905 | 5336 |
| MZ11324 | Pmg | M | 10/22/16 | 0 | Menominee | 45.5,-87.41 | 53.00 | 350 | 1.00 | 119650 | 48721 | 40.70 | 29456 | 160 |
| MZ11325 | Pmg | F | 10/23/16 | 0 | Menominee | 45.5,-87.41 | 42.90 | 350 | 1.00 | 269511 | 108061 | 40.10 | 46106 | 1688 |
| MZ11326 | Pmg | F | 10/23/16 | 0 | Menominee | 45.5,-87.41 | 44.60 | 350 | 1.00 | 79037 | 24894 | 31.50 | 17904 | 4 |
| MZ11327 | Pmg | M | 10/23/16 | 0 | Menominee | 45.5,-87.41 | 27.90 | 350 | 1.00 | 89295 | 31376 | 35.10 | 19566 | 103 |
| MZ11328 | Pmg | M | 10/23/16 | 0 | Menominee | 45.5,-87.41 | 154.00 | 350 | 1.00 | 2608601 | 980606 | 37.60 | 175557 | 73036 |
| MZ11329 | Pmg | M | 10/23/16 | 0 | Menominee | 45.5,-87.41 | 304.00 | 350 | 0.98 | 1508072 | 637781 | 42.30 | 94246 | 43490 |
| MZ11330 | Pmg | M | 10/23/16 | 0 | Menominee | 45.5,-87.41 | 58.00 | 350 | 0.99 | 114101 | 51363 | 45.00 | 19948 | 359 |
| MZ11331 | Pmg | M | 10/23/16 | 0 | Menominee | 45.5,-87.41 | 34.10 | 350 | 0.99 | 154619 | 78622 | 50.80 | 22912 | 4190 |
| MZ11332 | Pmg | F | 10/23/16 | 0 | Menominee | 45.5,-87.41 | 326.00 | 350 | 1.00 | 1867830 | 714744 | 38.30 | 150854 | 51548 |
| MZ11333 | Pmg | M | 10/23/16 | 0 | Menominee | 45.5,-87.41 | 31.10 | 350 | 0.99 | 170140 | 88778 | 52.20 | 33793 | 4935 |
| MZ176017 | Pln | F | 10/15/03 | 13 | Menominee | 45.51,-87.43 | 21.60 | 150 | 0.99 | 89863 | 22644 | 25.20 | 3162 | NA |
| MZ176018 | Pln | F | 10/15/03 | 13 | Menominee | 45.51,-87.43 | 11.70 | 150 | 0.98 | 192419 | 51019 | 26.50 | 10455 | NA |
| MZ176020 | Pln | F | 10/15/03 | 13 | Menominee | 45.5,-87.4 | 6.54 | 150 | 0.97 | 64924 | 24147 | 37.20 | 5603 | NA |
| MZ176021 | Pln | M | 10/15/03 | 13 | Menominee | 45.5,-87.4 | 12.70 | 150 | 0.97 | 134520 | 32733 | 24.30 | 7323 | NA |

**Table S2:** Specimen information and sequencing statistics for all specimens used in this study. Specimens beginning with MZ were sampled from the University of Michigan Museum of Zoology collection. The Age column indicates the specimen age at the time of DNA extraction (years old). Sequencing statistics indicate the concentration of DNA after extraction (Ext DNA conc), the starting amount of DNA used for library preparation (Starting DNA amt), the sample-specific bisulfite conversion rate measured using a spike-in of fully unmethylated phage DNA (Bisulfite conv rate), the number of read pairs retained after demultiplexing and cleaning (Demult read pairs), the number of read pairs retained after alignment (Aligned read pairs), the mapping efficiency (Mapping eff), and the total number of unique CpG positions sequenced at a read depth of 1X and 10X. The starred specimens were sequenced in two separate libraries and the reads from the two libraries were combined prior to methylation calling; therefore, the number of CpG positions reported in the last two columns represent this combined value.

| Specimen | Species | Sex | Date collected | Age (yo) | County | Location | Ext DNA conc (ng/$\mu$L) | Starting DNA amt (ng) | Bisulfite conv rate | Demult read pairs | Aligned read pairs | Mapping eff (%) | CpG pos (1X) | CpG pos (10X) |
|---|---|---|---|---|---|---|---|---|---|---|---|---|---|---|
| MZ176022 | Pln | M | 10/27/03 | 13 | Menominee | 45.5,-87.39 | 7.36 | 150 | 0.99 | 44797 | 7410 | 16.50 | 1565 | 2 |
| MZ176023 | Pln | M | 10/27/03 | 13 | Menominee | 45.5,-87.39 | 1.72 | 40 | 0.99 | 191600 | 35374 | 18.50 | 4441 | 2676 |
| MZ176024 | Pln | F | 10/27/03 | 13 | Menominee | 45.5,-87.39 | 13.40 | 150 | 0.99 | 954988 | 108678 | 11.40 | 14047 | 7 |
| MZ176026 | Pln | F | 10/28/03 | 13 | Menominee | 45.5,-87.39 | 40.80 | 150 | 0.99 | 325649 | 65356 | 20.10 | 10833 | 6 |
| MZ176027 | Pln | M | 10/28/03 | 13 | Menominee | 45.5,-87.39 | 4.64 | 150 | 1.00 | 83411 | 22934 | 27.50 | 3644 | NA |
| MZ176028 | Pln | F | 10/28/03 | 13 | Menominee | 45.5,-87.39 | 0.83 | 40 | 0.99 | 530725 | 351089 | 66.20 | 33067 | 26074 |
| MZ176029 | Pln | M | 10/29/03 | 13 | Menominee | 45.55,-87.33 | 5.28 | 150 | 0.98 | 132091 | 45778 | 34.70 | 7730 | NA |
| MZ176047 | Pln | M | 10/28/03 | 13 | Menominee | 45.5,-87.39 | 5.80 | 150 | 0.97 | 13742 | 649 | 4.70 | 183 | NA |
| MZ176048 | Pln | F | 10/28/03 | 13 | Menominee | 45.5,-87.39 | 21.60 | 150 | 0.98 | 213002 | 55793 | 26.20 | 7932 | NA |
| MZ176050 | Pln | F | 10/27/03 | 13 | Menominee | 45.5,-87.39 | 19.10 | 150 | 0.99 | 668606 | 134782 | 20.20 | 15395 | 4 |
| MZ176101 | Pmg | M | 10/27/03 | 13 | Menominee | 45.5,-87.39 | 7.41 | 150 | 0.98 | 35139 | 6399 | 18.20 | 1303 | 36 |
| MZ176107 | Pmg | M | 10/27/03 | 13 | Menominee | 45.5,-87.39 | 3.52 | 150 | 0.98 | 23853 | 5493 | 23.00 | 1013 | 3 |
| MZ176109 | Pmg | F | 10/27/03 | 13 | Menominee | 45.5,-87.39 | 1.75 | 40 | 0.99 | 59100 | 15227 | 25.80 | 1570 | 879 |
| MZ176111 | Pmg | F | 10/27/03 | 13 | Menominee | 45.5,-87.39 | 26.00 | 150 | 1.00 | 95420 | 20439 | 21.40 | 5174 | NA |
| MZ176115 | Pmg | M | 10/27/03 | 13 | Menominee | 45.5,-87.39 | 0.87 | 40 | 1.00 | 13468 | 94 | 0.70 | 24 | NA |
| MZ176116 | Pmg | F | 10/27/03 | 13 | Menominee | 45.5,-87.39 | 6.33 | 150 | 0.99 | 98053 | 7178 | 7.30 | 1846 | 18 |
| MZ176117 | Pmg | F | 10/27/03 | 13 | Menominee | 45.5,-87.39 | 1.27 | 40 | 0.99 | 13189 | 419 | 3.20 | 175 | 11 |
| MZ176123 | Pmg | M | 10/27/03 | 13 | Menominee | 45.5,-87.39 | 31.80 | 150 | 1.00 | 47125 | 12431 | 26.40 | 2726 | 2 |
| MZ176125 | Pmg | M | 10/27/03 | 13 | Menominee | 45.5,-87.39 | 37.50 | 150 | 0.98 | 90963 | 11079 | 12.20 | 2830 | 13 |
| MZ176127 | Pmg | M | 10/27/03 | 13 | Menominee | 45.5,-87.39 | 10.90 | 150 | 1.00 | 381001 | 70084 | 18.40 | 1663 | NA |
| MZ176129 | Pmg | F | 10/27/03 | 13 | Menominee | 45.5,-87.39 | 40.40 | 150 | 0.98 | 128604 | 16942 | 13.20 | 4338 | 2 |
| MZ176136 | Pmg | M | 10/27/03 | 13 | Menominee | 45.5,-87.39 | 1.74 | 40 | 1.00 | 7880 | 3804 | 48.30 | 246 | 171 |
| MZ176140 | Pmg | F | 10/27/03 | 13 | Menominee | 45.5,-87.39 | 15.60 | 150 | 0.99 | 49075 | 11947 | 24.30 | 1963 | 19 |
| MZ84532 | Pmg | M | 8/19/40 | 76 | Menominee | 45.41,-87.46 | 1.09 | 40 | 0.99 | 20561 | 3277 | 15.90 | 360 | 236 |
| MZ84533 | Pmg | M | 8/20/40 | 76 | Menominee | 45.41,-87.46 | 1.86 | 40 | 0.99 | 7083 | 68 | 1.00 | 39 | NA |
| MZ84534 | Pmg | F | 8/21/40 | 76 | Menominee | 45.41,-87.46 | 1.88 | 40 | 0.99 | 47104 | 175 | 0.40 | 74 | NA |
| MZ84535 | Pmg | F | 8/21/40 | 76 | Menominee | 45.41,-87.46 | 1.08 | 40 | 0.99 | 171764 | 20054 | 11.70 | 1231 | 787 |
| MZ84536 | Pmg | M | 8/21/40 | 76 | Menominee | 45.41,-87.46 | 2.90 | 40 | 0.99 | 52603 | 7738 | 14.70 | 3775 | 235 |
| MZ84537 | Pmg | M | 8/21/40 | 76 | Menominee | 45.41,-87.46 | 4.74 | 40 | 0.97 | 8746 | 1071 | 12.20 | 60 | 30 |
| MZ84540 | Pmg | M | 8/22/40 | 76 | Menominee | 45.41,-87.46 | 4.84 | 40 | 1.00 | 9081 | 736 | 8.10 | 130 | 18 |
| MZ84541 | Pmg | F | 8/22/40 | 76 | Menominee | 45.41,-87.46 | 2.38 | 40 | 0.98 | 1863 | 324 | 17.40 | 92 | 6 |

**Table S2:** Specimen information and sequencing statistics for all specimens used in this study. Specimens beginning with MZ were sampled from the University of Michigan Museum of Zoology collection. The Age column indicates the specimen age at the time of DNA extraction (years old). Sequencing statistics indicate the concentration of DNA after extraction (Ext DNA conc), the starting amount of DNA used for library preparation (Starting DNA amt), the sample-specific bisulfite conversion rate measured using a spike-in of fully unmethylated phage DNA (Bisulfite conv rate), the number of read pairs retained after demultiplexing and cleaning (Demult read pairs), the number of read pairs retained after alignment (Aligned read pairs), the mapping efficiency (Mapping eff), and the total number of unique CpG positions sequenced at a read depth of 1X and 10X. The starred specimens were sequenced in two separate libraries and the reads from the two libraries were combined prior to methylation calling; therefore, the number of CpG positions reported in the last two columns represent this combined value.

| Specimen | Species | Sex | Date collected | Age (yo) | County | Location | Ext DNA conc (ng/$\mu$L) | Starting DNA amt (ng) | Bisulfite conv rate | Demult read pairs | Aligned read pairs | Mapping eff (%) | CpG pos (1X) | CpG pos (10X) |
|---|---|---|---|---|---|---|---|---|---|---|---|---|---|---|
| MZ84625 | Pln | M | 8/17/40 | 76 | Menominee | 45.18,-87.62 | 3.96 | 40 | 0.99 | 17191 | 352 | 2.00 | 122 | 7 |
| MZ84627 | Pln | F | 8/18/40 | 76 | Menominee | 45.41,-87.46 | 1.42 | 40 | 0.96 | 13109 | 657 | 5.00 | 11 | 9 |
| MZ84628 | Pln | F | 8/19/40 | 76 | Menominee | 45.41,-87.46 | 1.96 | 40 | 0.99 | 28681 | 972 | 3.40 | 304 | 12 |
| MZ84629 | Pln | M | 8/19/40 | 76 | Menominee | 45.41,-87.46 | 1.94 | 40 | 0.99 | 4150 | 36 | 0.90 | 17 | NA |
| MZ84631 | Pln | F | 8/19/40 | 76 | Menominee | 45.41,-87.46 | 3.06 | 40 | 0.99 | 7106 | 176 | 2.50 | 48 | 4 |
| MZ84634 | Pln | M | 8/19/40 | 76 | Menominee | 45.41,-87.46 | 1.29 | 40 | 0.99 | 49897 | 1016 | 2.00 | 248 | 14 |
| MZ84636 | Pln | F | 8/20/40 | 76 | Menominee | 45.41,-87.46 | 1.20 | 40 | 0.94 | 3063 | 598 | 19.50 | 365 | NA |
| MZ84637 | Pln | F | 8/21/40 | 76 | Menominee | 45.41,-87.46 | 1.43 | 40 | 0.99 | 16009 | 2132 | 13.30 | 152 | 99 |
| MZ84639 | Pln | M | 8/21/40 | 76 | Menominee | 45.41,-87.46 | 1.33 | 40 | 1.00 | 7673 | 76 | 1.00 | 36 | NA |
| MZ84641HI* | *P. leucopus* | F | 8/21/40 | 76 | Menominee | 45.41,-87.46 | 9.64 | 150 | 0.99 | 6388 | 707 | 11.10 | 6729 | 5005 |
| MZ84641LO* | see above | - | - | - | - | - | - | 40 | 0.98 | 281116 | 87695 | 31.20 | - | - |
| MZ84642 | Pln | F | 8/21/40 | 76 | Menominee | 45.41,-87.46 | 3.24 | 40 | 0.98 | 8917 | 198 | 2.20 | 54 | 2 |
| MZ84645 | Pln | M | 8/21/40 | 76 | Menominee | 45.41,-87.46 | 0.68 | 40 | 0.99 | 15964 | 2443 | 15.30 | 310 | 99 |
| MZ84657HI* | *P. leucopus* | M | 8/26/40 | 76 | Menominee | 45.41,-87.46 | 48.60 | 150 | 0.99 | 582375 | 189843 | 46.20 | 18318 | 212 |
| MZ84657LO* | see above | - | - | - | - | - | - | 40 | 0.99 | 11410 | 5267 | 32.60 | - | - |
