## Supplemental Methods for "Museum epigenomics: characterizing cytosine methylation in historic museum specimens"

This document contains specialized methods for library preparation using historic bone samples. Included are: a protocol for sampling microturbinates, two DNA extraction protocols (a phenol-chloroform protocol and a modified Qiagen DNeasy extraction protocol), and a bisulfite ddRAD protocol for describing DNA methylation. All work performed prior to barcode ligation should be performed in a dedicated laboratory space following specialized protocols for working with ancient or historic DNA.

### Microturbinate sampling protocol:

Microturbinates (small nasal bones) can be sampled as a minimal damage option for sampling skull tissue (**??**). Homogenizing the tissue prior to DNA extraction may help increase yield.

1. Sterilize instruments with 10% bleach, followed by 95% ethanol.
2. Tare a piece of sterile foil on scale.
3. Insert narrow micropick into nasal cavity through right nostril. Gently scrape around outside edge.
4. Tilt skull to allow material to fall out of nostril. Tap gently to help dislodge.
5. Repeat until collect 4 - 11 mg of tissue.
6. Transfer bone tissue from foil into sterile 2 mL thick-walled microcentrifuge tube (compatible with tissue homogenizer).

Perform the following steps immediately before beginning the extraction:

1. Place 4 sterile 2.4 mm stainless steel beads into each tube of bone tissue.
2. Homogenize in FastPrep tissue homogenizer for 1 minute at 6.0 m/s. Repeat.

### Phenol Chloroform Extraction:

Phenol chloroform extraction is recommended for older historic specimens; for less degraded specimens, the following Qiagen protocol may be followed instead. Phase lock gel tubes are not required but may improve the yield of extracted DNA by making it easier to decant the full volume at each step. Amicon filters were used to purify the DNA, but alternative filters or ethanol precipitation could be used at this stage instead. Extracted DNA should be stored frozen in small aliquots to reduce the number of freeze-thaw cycles. This protocol is modified from **?** and the University of Michigan Genomic Diversity Lab phenol-chloroform protocol. It has been altered for a lower volume reaction.

**Reagents**

1. Chelation
   (a) EDTA 0.5 M pH 8 (1000 $\mu$L per sample)
2. Digestion
   (a) 15 mM Tris-Hcl pH 8 (140 $\mu$L per sample)
   (b) SDS 10 % w/v (20 $\mu$L per sample)
   (c) Proteinase K, 25 mg/mL (20 $\mu$L per sample)
   (d) DTT, 500 mM (20 $\mu$L per sample)
3. Phase separation
   (a) Phenol pH 6.6 (400 $\mu$L per sample)
   (b) Chloroform (200 $\mu$L per sample)
4. Elution

(a) Amicon Ultra 0.5 mL filters (30kDa)

**Day 1**

1. Add 1000 $\mu$L of EDTA 0.5 M pH 8 to each sample. Mix by inversion.
2. Incubate on rotator overnight at room temperature.

**Day 2**

1. Centrifuge at 4000 x g for 10 min until bone forms pellet at bottom tube.
2. Remove eluate.
3. Add 140 $\mu$L of 15 mM Tris-Hcl pH 8 to tube containing bone pellet.
4. Add 20 $\mu$L SDS.
5. Add 20 $\mu$L proteinase K.
6. Add 20 $\mu$L DTT. Incubate on rotator overnight at 55° C.

**Day 3**

1. Prepare two 2 mL screw top tubes per sample containing 200 $\mu$L phenol pH 6.6 and one 2 mL screw top tube per sample containing 200 $\mu$L chloroform.
2. Spin digested sample briefly, add 100 $\mu$L molecular grade water, and add to phenol tube 1.
3. Rotate at room temperature for 10 min.
4. Spin gel tubes at 16,000 x g for 20-30 sec.
5. Add to phase lock gel tube and shake vigorously for 15 sec or until homogenous (do not vortex).
6. Centrifuge at 16,000 x g for 5 min.
7. Decant aqueous layer (top) into phenol tube 2.
8. Rotate at room temperature for 10 min.
9. Add to phase lock gel tube and shake vigorously for 15 sec or until homogenous (do not vortex).
10. Centrifuge at 16,000 x g for 5 min.
11. Decant aqueous layer (top) into chloroform tube.
12. Rotate at room temperature for 5 min.
13. Add to phase lock gel tube and shake vigorously for 15 sec or until homogenous (do not vortex).
14. Centrifuge at 16,000 x g for 5 min.
15. Moisten Amicon filters with small amount of molecular grade water, then transfer aqueous layer into same column.
16. Spin at 8,000 x g for 5 minutes until sample has completely passed through membrane. Discard filtrate.
17. Add 450 $\mu$L molecular grade water and spin again at 8,000 x g for 3 minutes (or until final volume $< 100\mu$L).
18. Reverse spin: Place device upside down in new microcentrifuge tube and spin for 2 minutes at 1,000 x g.
19. Estimate volume and reconstitute with water.

### Modified Qiagen DNeasy Extraction:

This protocol is modified from the Qiagen DNeasy Blood and Tissue Kit protocol with added steps for increasing yield. Specifically, we recommend increasing lysis incubation times, increasing the volume of proteinase K, and increasing the elution incubation time and temperature. To increase the concentration of the final eluted sample, we recommend reducing the elution volume and keeping the first and second elutions separate. Also see **???**.

1. Add 180 µL Buffer ATL and 20 µL proteinase K to sample, vortex on low.
2. Incubate at 56°C for 48 hours, or until fully lysed (no solids visible). During incubation, add 20 µL fresh proteinase K after 24 hours.
3. Vortex for 15 s. Add 200 µL Buffer AL, vortex thoroughly, incubate at 70°C for 10 min. Add 200 µL 100% ethanol and vortex throughly.
4. Pipet full mixture into spin column, spin at 6000 x g (8000 rpm) for 1 min. Discard flow through.
5. Place column in new tube, add 500 µL Buffer AW1, spin for 1 min at 6000 x g (8000 rpm). Discard flow through.
6. Place column in new tube, add 500 µL Buffer AW2, spin for 3 min at 20,000 x g (14,000 rpm) to dry membrane. Carefully remove collection tube and discard flow through.
7. Warm molecular grade water to 70°C. Place column in 1.5 mL tube. Pipet 80 µL (or desired elution volume) warmed water directly onto membrane, let sit at room temperature for 5 minutes. Spin for 1 min at 6000 x g (8000 rpm).
8. Repeat elution step.

**Bisulfite ddRAD Library Preparation:**

This protocol uses a combination of ddRAD and bisulfite treatment to produce reduced representation methylomes. This protocol requires methylated adapters - these were custom ordered from Sigma Aldrich. These libraries were prepared using the restriction enzymes Sph-HF and MluCI, however, other enzyme pairs could easily be substituted. This protocol is for combinatorial multiplexing with P1 adapter barcodes and PCR primer indices. Here we size selected for fragments 325-425 bp in length; targeting shorter fragments might be preferable for older or more degraded samples. We recommend including a spike-in of fully unmethylated lambda phage DNA to directly measure the efficiency of the bisulfite conversion reaction. For older or degraded specimens, we recommend using a low concentration for the spike-in - here we used 0.1% of the sample concentration. See **?** for more details about the ddRAD protocol, including adapter sequences, and **??** for similar bisulfite RAD protocols.

**DNA preparation:**

1. Prepare 17 $\mu$L of each sample at a standardized DNA concentration (500 - 1000 ng), place in separate 0.2 mL tubes. Include one sample of fully unmethylated lambda phage DNA.

**Digest:**

1. Prepare master mix: per sample, 2 $\mu$L 10X CutSmart buffer, 0.25 $\mu$L Sph-HF, 0.5 $\mu$L MluCI, 0.25 $\mu$L water. Add enzymes last, mix by pipetting (do not vortex). Keep all reagents ice cold during plate preparation.
2. Add 3 $\mu$L master mix to 17 $\mu$L DNA samples.
3. Digest for 1 hour at 37°C. Cool reaction to room temperature.
4. Clean digested DNA (e.g., Ampure cleanup) and reconstitute in 33 $\mu$L.
5. Quantify concentrations of cleaned digests and digested phage.
6. Create new plate of standardized digests in 32 $\mu$L. Dilute lambda phage DNA to desired spike-in concentration (in this protocol 0.1% of sample concentration) in 1 $\mu$L and add to digests for total volume of 33 $\mu$L.

**Adapter ligation:**

1. Prepare working strength annealed adapters as described in **?**. Keep all reagents ice cold.

2. Prepare master mix of 1 $\mu$L adapter P2 working stock, 4 $\mu$L 10X ligase buffer, 1 $\mu$L T4 ligase.
3. Add 1 $\mu$L adapter P1 working stock to standardized digest plate.
4. Add 6 $\mu$L master mix to each plate (total reaction volume is 40 $\mu$L).
5. Incubate at room temperature for 30 min, then heat-kill at 65°C for 10 min. After the heat-kill, cool the solution at 2°C per 90 seconds until it reaches room temp.
6. Pool into 10 sublibraries of uniquely barcoded individuals.
7. Clean pooled sublibraries (e.g., Ampure cleanup) and reconstitute in 30 $\mu$L.

**Size selection with Sage Science Pippin-Prep:**

1. Run size selection on 30 $\mu$L of pooled sublibraries following Pippin Prep instructions for appropriate range size.
   (a) This study: 376-412 bp for high concentration library, 325-425 bp for medium and low concentration libraries.
2. Recover sample from elution well.

**Bisulfite Conversion:**

(Using Promega MethylEdge kit - several similar commercial kits are available.)

1. Place 20 $\mu$L DNA into a 0.2 mL PCR tube.
2. Add 130 $\mu$L Bisulfite ME Conversion Reagent to each sample, pipette gently to mix. Spin briefly to collect sample at bottom of tube.
3. Incubate in thermocycler.
   (a) 8 minutes at 98°C
   (b) 60 minutes at 54°C
   (c) Hold at 4°C

DNA Desulphonation and Cleanup

1. Place one ME Spin Column per sample into a collection tube.
2. Add 600 $\mu$L ME Binding Buffer to spin column. Transfer entire bisulfite-treated sample to the column. Close the cap and mix by inverting tube several times.
3. Spin at maximum speed for 30 s. Remove filter, discard flow through, place filter back on same collection tube.
4. Add 100 $\mu$L 1X ME Wash Buffer. Spin at maximum speed for 30 s. Remove filter, discard flow through, place filter back on same collection tube.
5. Add 200 $\mu$L ME Desulphonation Buffer. Close caps and incubate at room temperature for 15 min.
6. Spin at max speed for 30 s. Remove filter, discard flow through, place filter back on same collection tube.
7. Add 200 $\mu$L ME Wash Buffer. Spin at max speed for 30 s.
8. Repeat wash step.
9. Place spin column onto a clean 1.5 mL microcentrifuge tube.
10. Add 20 $\mu$L ME Elution Buffer and spin at max speed for 30 s.
11. Remove and discard spin column.

**PCR:** Set up reaction on ice.

Total reaction:
KAPA HiFi HotStart Uracil+ ReadyMix (2X): 25 $\mu$L
Universal Primer 1 (10X): 2.5 $\mu$L
Barcoded Primer 2 (10X): 2.5 $\mu$L
DNA: 20 $\mu$L
**Total: 50 $\mu$L**

1. Prepare Master Mix:
   (a) KAPA HiFi HotStart Uracil+ ReadyMix (2X): 25 $\mu$L

   (b) Universal Primer 1 (10X): 2.5 $\mu$L
2. Add 2.5 $\mu$L barcoded Primer 2 (10X) to 20 $\mu$L DNA.
3. Add 27.5 $\mu$L master mix.
4. Mix thoroughly by pipetting, spin briefly.
5. Amplify in thermocycler.
   (a) all other sublibraries
      i. 98°C 45 s
      ii. 98°C 15 s |
      iii. 60°C 30 s |     X 17
      iv. 72°C 30 s |
      v. 72°C 1 m
6. Clean amplified sublibraries (e.g., Ampure cleanup) and elute to desired volume.
7. Quantify sublibraries, standardize concentrations, and pool into final library for sequencing.
